## Supplementary material for "Hippocampal and medial prefrontal cortices encode structural task representations following progressive and interleaved training schedules": S1 Text

### Interleaved and progressive training in multilayer perceptrons

#### Results and Discussion

Here we demonstrate that different training schedules can guide even very simple agents to learn fundamentally different task representations which, in turn, affects generalisation. Across multiple independent runs, we trained multilayer perceptrons (MLPs) with 6 different architectures to perform a set of binary discriminations similar in structure to our fMRI task (see Figure 1B). These discriminations were presented either progressively (one after another), or in an interleaved order (see Figures 1C and 1D). We then tested the perceptrons on both explicitly trained (premise) and transitively inferred discriminations.

The goal of this analysis was to characterise effects of each training schedule on learning and inference. As such, the MLPs were designed to be as simple as possible while having the ability to learn a task representation that could support transitive inference. They were not indented to be a faithful model of how humans solve such tasks, nor is this analysis indented to adjudicate between retrieval- and encoding-based models of generalisation. The MLPs consisted of three layers (input, hidden, and output) with varying numbers of hidden units (between 1 and 6) connected to all inputs and output (see Figure S1.1A). In short, training involved presenting the perceptrons with two input symbols (e.g., 'A' and 'B') coded as a binary vector and updating the network weights via backpropagation to reproduce the "correct" symbol as a one-hot vector on the output layer.

We explicitly trained 6 premise discriminations; 'A>B', 'B>C' ... 'F>G' (correct responses indicated to the left of the greater-than sign) over 3,600 training steps (note: this is 10x the number of training trials given to human participants in preparation for the fMRI task, see methods section below). As such, these contingencies implied a 1D transitive hierarchy (A>B>C>D>E>F>G). For the networks with only one hidden unit, the output layer only received a 1-dimensional latent representation of the inputs and so high-levels of performance depended on non-trivial weights and bias distributed between layers. Nonetheless, many solutions that do yield near-perfect performance on both premise and inferred discriminations do exist (see https://osf.io/ps3ch/ for a scripted example). With more hidden units the networks were capable of learning increasingly sparse representations of each discrimination, see [1,2]. For example, with 6 hidden units, a specific hidden unit could uniquely activate the target output for each trained discrimination.

Interleaved training involved presenting the MLPs with all 6 discriminations in a pseudorandom order such that there was a uniform probability (1/6) of encountering any one on a particular trial (see Figure 1C). As such, training-related prediction error (as measured by a cross-entropy cost function) gradually decreased across interleaved training trials (see Figure S1.1B). In contrast, progressive training involved 6 epochs of different lengths that gradually introduced discriminations and ensured that, once introduced, they were presented in all subsequent epochs. Given this, progressive training was characterised by sharp increases, and then rapid decreases in prediction error at the start of each epoch. For each MLP architecture, interleaved and progressive training runs were repeated 10,000 times with varied training orders and different network initialisations (i.e., random starting parameters).

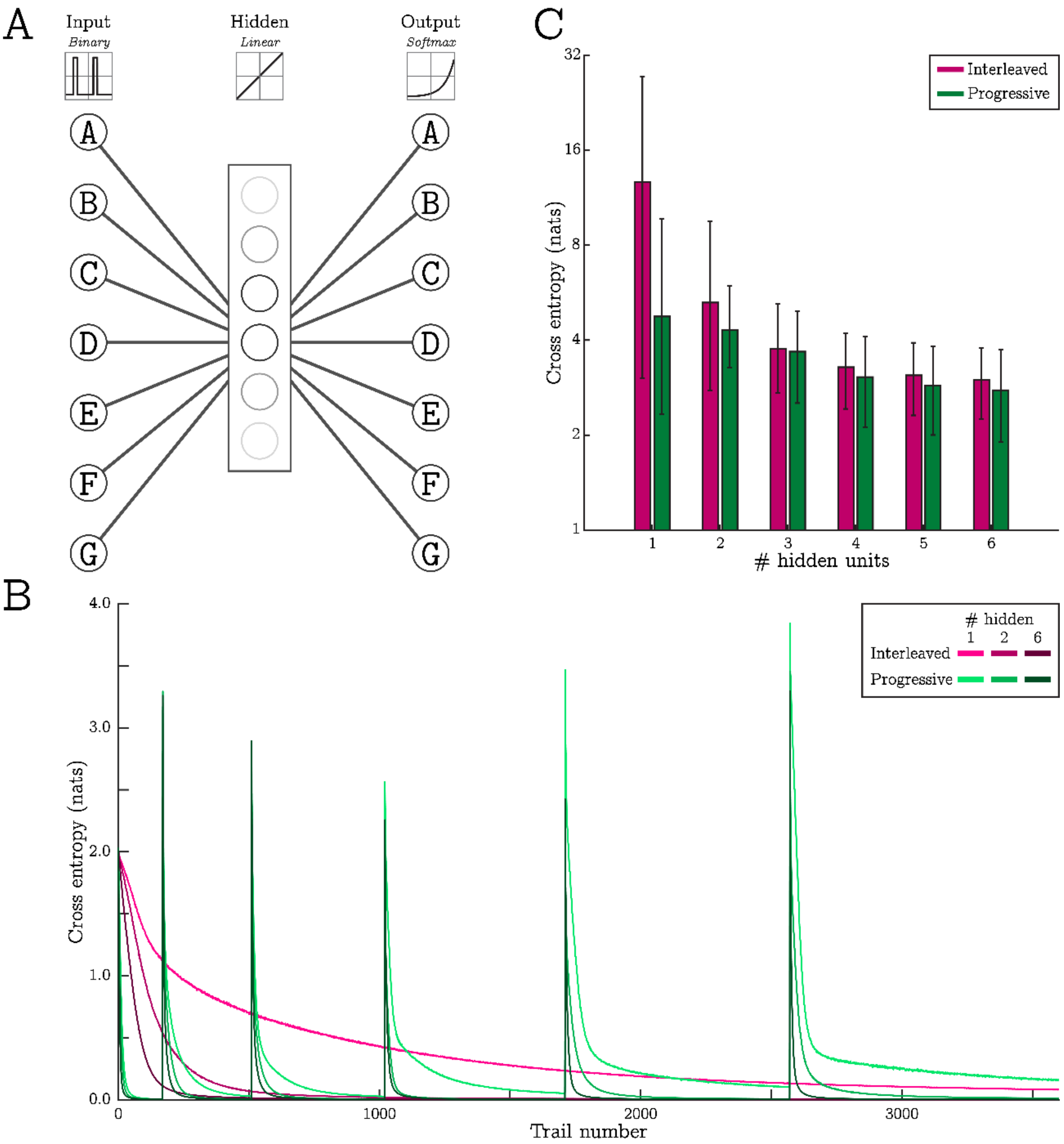


Figure S1.1 Multilayer perceptrons (MLPs) show better generalisation in a transitive interference task after progressive training, particularly when the hidden layer only contains a single unit. A) An illustration of the 3-layer MLPs included in the analysis. Seven input units (labelled A-G, one for each discriminable feature) provided weighted binary inputs to between 1 and 6 biased hidden units with a linear activation function. In turn, activity in the hidden units projected along weighted connections to seven biased outputs with a SoftMax activation function. B) We trained the MLPs on the six premise discriminations in Figure 1 via backpropagation. Training was structured in two ways (see Figures 1C and 1D). With interleaved training, a cross entropy loss function decreased exponentially over trials (magenta lines) whereas progressive training resulted in rapid exponential decreases following the start of each epoch (green lines). Note: the learning curves plotted in this figure result from averaging 10,000 independent training runs for networks with either 1, 2 or 6 hidden units. C) After each training run, we tested the performance of MLPs on the full set of inferred discriminations show in Figure 1. Inference performance as measured by the cross-entropy loss improved as more hidden units were added and was invariably better following progressive training. Nonetheless, the effect of progressive training was most pronounced for MLPs with a single hidden unit. Bars indicate mean cross-entropy scores across runs on a log-scale, error bar ranges include 95% of scores.

We tested the performance of MLPs in two ways: *1)* using the same cross-entropy measure employed during training and, *2)* using a 2-alternative Luce choice (2-ALC) decision rule implemented by a SoftMax normaliser (see Supplementary Methods). Unlike cross-entropy, the 2-ALC measure allowed us to examine whether the activity of the target output unit was greater than the activity of the concurrently presented non-target (which tended to be highly activated by its corresponding input). Both interleaved and progressive training schedules resulted in all MLPs learning the 6 premise discriminations to a high level (Table S1.1). According to both performance measures, premise performance increased as hidden units were added, and were slightly better following interleaved training.

We then tested the MLP on 6 transitive inference trials that were not explicitly trained (e.g., 'B>D'). Cross-entropy statistics indicated that progressively trained MLPs invariably performed better on these inference trials than MLPs trained via an interleaved schedule (Figure S1.1C). In networks with a single hidden unit, interleaved training did not generally promote greater activity in the target output relative to the non-target output, 2-ALC accuracy: 49.8% (CI: +/- 0.367; see Figure S1.2A). However, the variability in inference performance across different interleaved training runs was large (95% range: 17.0 - 78.8%). In contrast, progressive training in these single hidden unit networks lead to substantially higher levels of inferential accuracy: 64.1% (CI: +/- 0.174), yet the variability in performance reduced (95% range: 51.6 - 83.7%). This pattern in 2-ALC scores did not hold for networks with two or more hidden units, which tended to activate non-target output units more strongly following both training schedules (Table S1.2).

The above demonstrates that progressive training can yield better inference in some MLPs, particularly those with only a single hidden unit. To explore the reasons for this, we computed the linear activation of each output unit (prior to SoftMax normalisation) that resulted from stimulating the corresponding input unit with a one-hot vector. We denote this activation, $\Delta\alpha_i^3 \mid \alpha_i^1 = 1$, for the $i^{\text{th}}$ input/output unit, where superscripts denote layer indices. Figure S1.2B plots these values as a function of training trial, averaged across runs for MLPs with a single hidden unit. This showed that activity in the output corresponding to stimulus 'A' is generally higher than the activity of all others, activity in 'B' is next highest, and so on. As such, progressive training produces activations in each output unit that faithfully represents a generalised value for the corresponding stimulus as long as that stimulus is presented on the input layer. We note that this is also evident in MLPs with more than one hidden unit yet, the value representation is much weaker (see https://osf.io/2fw3g/).

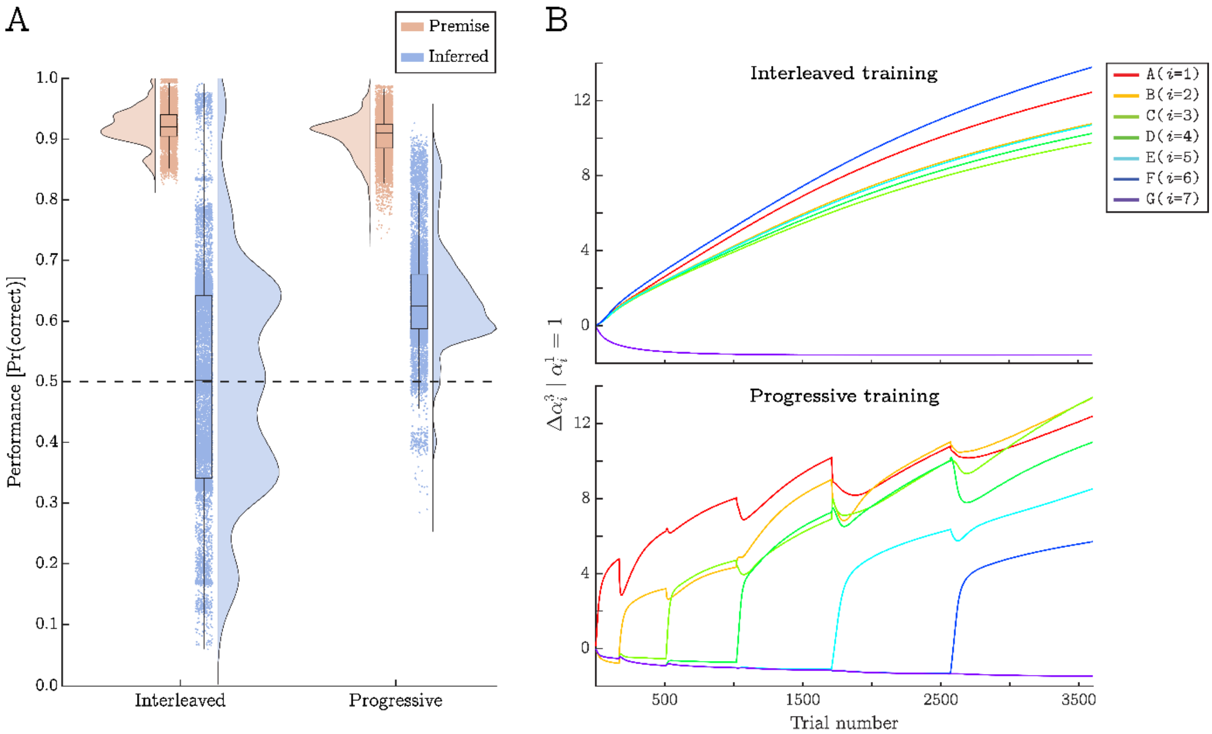


Figure S1.2 Progressive training promotes transitive inference in MLPs with a single hidden unit due to the learning of a generalised value gradient. A) We tested the performance of the MLP on both trained and inferred discriminations using a SoftMax function to implement a 2-alternative Luce choice decision (see *Eq. S1.1*). Performance on the premise discriminations was comparable across training methods but average performance on inference trials was much higher following progressive training. Despite this, the variability in inference performance that resulted from interleaved learning was large such that some MLP models trained in this way achieved very high levels of performance. B) When an input stimulus is presented as a one-hot vector, activity in corresponding output unit reflects a generalised value gradient following progressive, but not interleaved, training. Different lines plot activations in different output units (averaged across training runs). With an interleaved training procedure, activity levels are approximately equivalent for all stimuli that are rewarded during training. During progressive training, activity in the output corresponding to stimulus ‘A’ is generally higher than the activity of all others, activity in ‘B’ is next highest, and so on. This happens because learning the first discrimination (‘A>B’) tunes the MLP to increase the activation of the ‘A’ output above all others. When the next discrimination is introduced (‘B>C’), activity in the ‘B’ output starts to increase yet, because the ‘A>B’ discrimination is still being tested, activation in the ‘A’ output also must rise to maintain performance. This pattern continues down the transitive hierarchy as long as older discriminations are presented often enough to avoid catastrophic interference.

This generalised value gradient develops because learning the first discrimination (‘A>B’) tunes the MLP to increase the activation of the ‘A’ output above all others. When the next discrimination is introduced (‘B>C’), activity in the ‘B’ output starts to increase yet, because the ‘A>B’ discrimination is still being tested, activation in the ‘A’ output also must rise to maintain performance. This pattern continues down the transitive hierarchy as long as older discriminations are presented often enough to avoid catastrophic interference. As such, progressive training can yield a very specific error gradient that allows the network to learn a generalised value for each stimulus. The same is not true for MLPs trained via an interleaved schedule. Here, activity levels in output units are approximately uniform for all stimuli that are rewarded during training (i.e., excluding stimulus 'G'). Instead, networks trained in this way tended to learn the discriminations by inhibiting the activity of non-target outputs in response to the presentation of specific premise pairs.

A consequence of progressive training is that it biased MLPs with a single hidden unit towards learning a limited number of solutions to the discrimination problem (see Figure S1.3). We used a gaussian mixture model to characterise the distribution of trained parameters and found that they tended towards 53 distinct 'attractors' after interleaved training, but only 28 attractors after progressive training. Summed variance scores across parameters revealed that interleaved training resulted in more diverse solutions relative to progressive training ($\sigma^2 =$ 328 and 177, respectively). This patten was also evident for MLPs with two or more hidden units, although the difference between training schedules was much less pronounced (see Table S1.3).

For the above analyses, all MLPs were trained with a learning rate of 0.1 such that sufficient levels of performance on the premise pairs were achieved after 3,600 trials (10 times the number in the pre-scanner task). However, this learning rate is relatively large comped to those used in most machine learning applications. As such, we ran a secondary analysis that involved training the MLPs with a learning rate of 0.01 over 36,000 trials. This only involved 1,000 independent runs per training schedule in order to limit computer memory usage. The secondary MLP analysis revealed a pattern of results that was identical to those reported above (see https://osf.io/udmsr/).

In sum, we find that progressively ordering contingencies in a transitive inference task can have a significant effect on the type of representations that are learned by MLPs, especially when they are constrained to rely on low-dimensional solutions (having only a single hidden unit). This happens because progressive training induces an error gradient that favours learning the relative value of all stimuli (A>B>C...). Generalisation performance increases as a direct result of this, yet progressive training also appears to reduce the diversity of learnt representations thereby precluding the development of models that achieve very high levels of performance. Progressive training does not confer the same effects in MLPs with two or more hidden units since these networks were biased toward learning localist representations of the premise pairs that did not encode the higher-order transitive structure.

Using similar MLPs, Flesch et al [1] found that networks initialised with high-variance parameters tended to learn 'lazy' representations of stimuli in a categorisation task that indiscriminately preserved irrelevant features. On the other hand, when the networks were initialised with low-variance parameters, they were biased towards learning low-dimensional (compressed) representations of the input space that linearly encoded task-relevant features. These highly structured, 'rich' representations were learnt slower than lazy ones but were more resilient noise in the input layer. Our findings extend these results by highlighting that progressive training may selectively promote generalisation when representations using rich coding schemes are acquired during training.

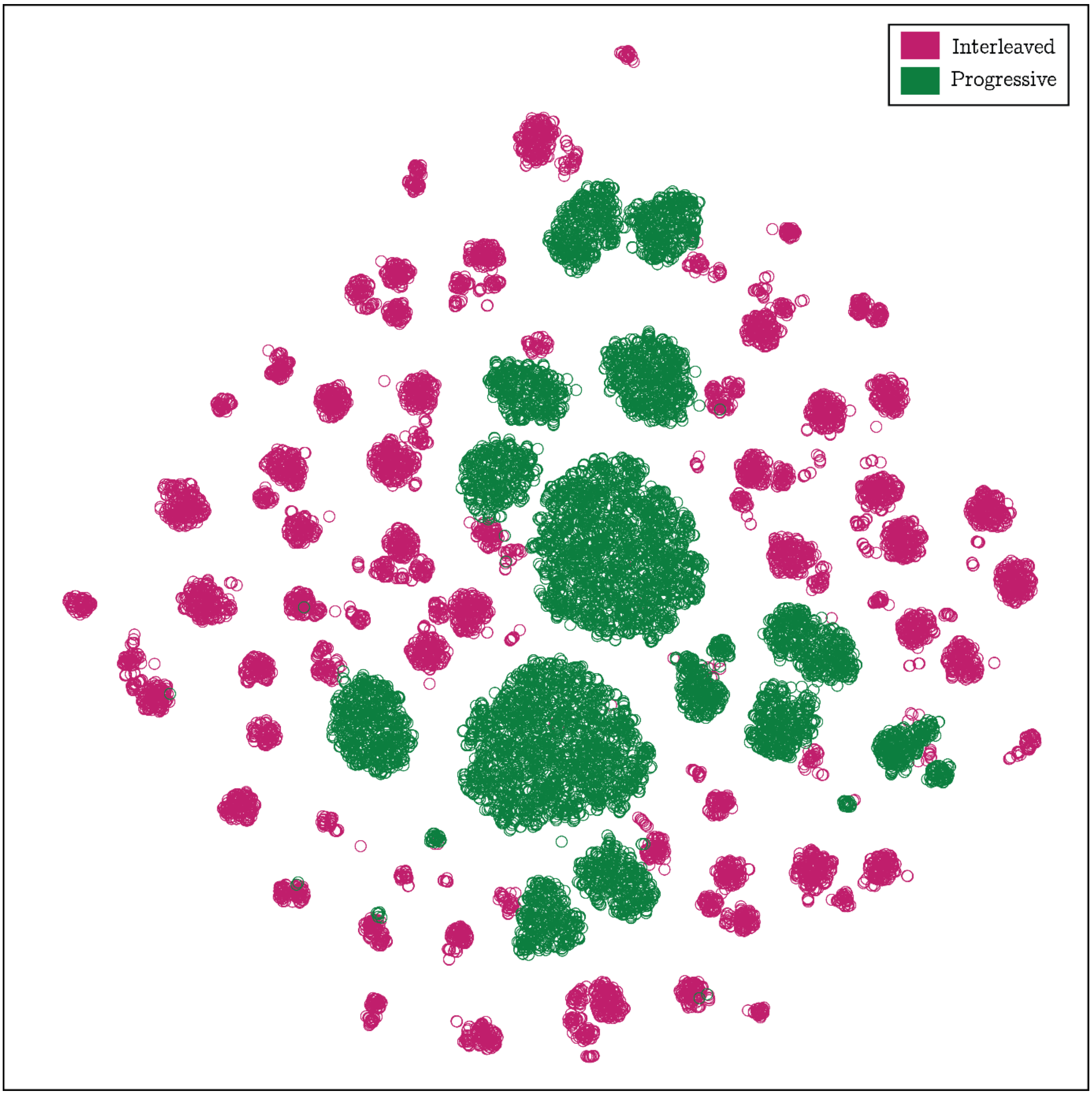


Figure S1.3 Progressive training in MLPs with a single hidden unit biases the networks towards learning a limited number of solutions. The plot shows a t-distributed stochastic neighbour embedding (tSNE) of the 22 network parameters generated in each training iteration (N=10,000 per training schedule). tSNE allows the visualisation many parameter-sets in a 2-dimentaional space by reducing dimensionality in a way that preserves the local structure within clusters of similar parameters. Independent of this visualisation, a gaussian mixture model detected the presence of 53 clusters following interleaved training, but only 28 clusters following progressive training. Summed variance scores across parameters revealed that interleaved training resulted in less clustered solutions relative to progressive training ($\sigma^2$ = 328 and 177, respectively).

Table S1.1 Performance of the MLPs on premise trials as reported in the main text. Cross-entropy (x-entropy) statistics are presented in natural units (nats), and 2-ALC accuracy scores are presented as proportions between 0 and 1. Rounded parentheses represent 95% confidence intervals and square braces indicate ranges that include 95% of scores.

| Number of hidden units | Interleaved training | | | Progressive training | |
|---|---|---|---|---|---|
| | *x-entropy* | *2-ALC* | | *x-entropy* | *2-ALC* |
| 1 | 0.081<br>(±$1.67 \times 10^{-4}$)<br>[0.072, 0.097] | .921<br>(±$5.70 \times 10^{-4}$)<br>[.864, .967] | | 0.335<br>(±$1.97 \times 10^{-3}$)<br>[0.224, 0.535] | .904<br>(±$6.49 \times 10^{-4}$)<br>[.844, .954] |
| 2 | 0.003<br>(±$2.31 \times 10^{-5}$)<br>[0.002, 0.005] | .977<br>(±$6.65 \times 10^{-5}$)<br>[.973, .984] | | 0.014<br>(±$8.42 \times 10^{-5}$)<br>[0.008, 0.021] | .964<br>(±$1.30 \times 10^{-4}$)<br>[.954, .976] |
| 3 | 0.001<br>(±$3.15 \times 10^{-6}$)<br>[0.001, 0.002] | .980<br>(±$4.36 \times 10^{-5}$)<br>[.977, .984] | | 0.005<br>(±$2.32 \times 10^{-5}$)<br>[0.003, 0.007] | .966<br>(±$7.40 \times 10^{-5}$)<br>[.961, .974] |
| 4 | 0.001<br>(±$1.92 \times 10^{-6}$)<br>[<0.001, 0.001] | .980<br>(±$2.56 \times 10^{-5}$)<br>[.978, .982] | | 0.003<br>(±$1.54 \times 10^{-5}$)<br>[0.002, 0.005] | .965<br>(±$5.06 \times 10^{-5}$)<br>[.961, .970] |
| 5 | 0.001<br>(±$1.48 \times 10^{-6}$)<br>[<0.001, 0.001] | .980<br>(±$1.97 \times 10^{-5}$)<br>[.978, .982] | | 0.003<br>(±$1.21 \times 10^{-5}$)<br>[0.002, 0.004] | .965<br>(±$4.25 \times 10^{-5}$)<br>[.961, .968] |
| 6 | 0.001<br>(±$1.22 \times 10^{-6}$)<br>[<0.001, 0.001] | .980<br>(±$1.65 \times 10^{-5}$)<br>[.979, .981] | | 0.003<br>(±$1.03 \times 10^{-5}$)<br>[0.002, 0.004] | .964<br>(±$3.84 \times 10^{-5}$)<br>[.961, .968] |

Table S1.2 Performance of the MLPs on inferred trials as reported in the main text. Cross-entropy (x-entropy) statistics are presented in natural units (nats), and 2-ALC accuracy scores are presented as proportions between 0 and 1. Rounded parentheses represent 95% confidence intervals and square braces indicate ranges that include 95% of scores.

| Number of hidden units | Interleaved training | | | Progressive training | |
|---|---|---|---|---|---|
| | *x-entropy* | *2-ALC* | | *x-entropy* | *2-ALC* |
| 1 | 12.7<br>(±0.142)<br>[3.03, 27.4] | .498<br>(±.004)<br>[.170, .788] | | 4.75<br>(±0.045)<br>[2.33, 9.69] | .641<br>(±.002)<br>[.516, .837] |
| 2 | 5.26<br>(±0.042)<br>[2.77, 9.54] | .513<br>(±.003)<br>[.270, .696] | | 4.30<br>(±0.018)<br>[2.28, 5.95] | .474<br>(±.001)<br>[.356, .572] |
| 3 | 3.760<br>(±0.016)<br>[2.72, 5.22] | .432<br>(±.002)<br>[.293, .607] | | 3.68<br>(±0.015)<br>[2.53, 4.95] | .356<br>(±.002)<br>[.244, .519] |
| 4 | 3.28<br>(±0.011)<br>[2.42, 4.21] | .398<br>(±.002)<br>[.283, .537] | | 3.05<br>(±0.012)<br>[2.12, 4.11] | .347<br>(±.001)<br>[.244, .465] |
| 5 | 3.09<br>(±0.010)<br>[2.31, 3.93] | .381<br>(±.001)<br>[.277, .502] | | 2.87<br>(±0.011)<br>[2.00, 3.82] | .342<br>(±.001)<br>[.241, .453] |
| 6 | 2.99<br>(±0.009)<br>[2.25, 3.78] | .372<br>(±.001)<br>[.275, .482] | | 2.78<br>(±0.011)<br>[1.90, 3.74] | .339<br>(±.001)<br>[.238, .449] |

Table S1.3 Variance in MLP parameters over 10,000 independent training runs, summed across all the parameters in each network (note: MLPs with more hidden units had more parameters). Progressive training results in less variable parameters, yet this effect is minimal for MLPs with three or more hidden units.

| Number of hidden units | Interleaved training | Progressive training |
|---|---|---|
| 1 | 328 | 177 |
| 2 | 119 | 100 |
| 3 | 89.4 | 88.5 |
| 4 | 85.1 | 83.8 |
| 5 | 84.0 | 82.7 |
| 6 | 83.6 | 82.3 |

#### Methods

As noted, we examined interleaved and progressive training in 6 different multilayer perceptrons (MLPs) with similar architectures. Each MLP consisted of 3 fully connected layers: 1) an input layer with 7 binary inputs, 2) a linearly activated hidden layer with between 1 and 6 biased hidden units, and 3) a SoftMax output layer with 7 biased outputs (see Figure S1.1). These architectures were chosen to be as simple as possible while still having the ability to learn parameters that would support near-perfect performance on both premise and inferred discriminations (see https://osf.io/ps3ch/).

Before training, the network was initialised with uniformly distributed random weights and biases sampled from the interval [-0.5, 0.5]. During training, we presented the network with 6 premise discriminations that had the same structure as those in the main behavioural task (i.e., A>B, B>C, etc.). This was done by setting the activity of two input units (e.g., A & B) to a value of 1 and setting all other input units (e.g., C, D, E, F, & G) to a value of 0. After each forward pass, a cross-entropy cost-function was used to quantify the error between the actual and target activity patterns on the output layer. The target patterns respected the contingencies described previously such that, when (e.g.) 'F>G' was presented on the input, output 'F' should have been maximally active relative to all other outputs. Backpropagation was then used to adjust all network weights and biases with a learning rate of 0.1. This rate parameter was chosen to allow for sufficient levels of learning following 3,600 training trials (10 times the number used in the pre-scanner task, see https://osf.io/uzyb7/). However, to examine the sensitivity of our results to this rate, we ran a secondary analysis that involved training the MLPs with a learning rate of 0.01 over 36,000 trials. This yielded an identical pattern of results to those presented above (see https://osf.io/udmsr/).

The MLPs were trained using both interleaved and progressive schedules across many independent iterations. As before, interleaved training involved presenting all 6 discriminations in a pseudorandom order such that there was a uniform probability of each discrimination occurring on any particular trial (10,000 iterations per MLP architecture). Progressive training involved 6 sequential epochs of different lengths which gradually introduced each discrimination one-by-one (10,000 iterations per architecture). The only difference between the training schedules used in the behavioural task and those used to train the MLP was the number of trials.

After training, we tested the MLP on the premise and inferred discriminations detailed in Figure 1B, quantifying performance using both the cross-entropy cost function and 2-alternative Luce choice (2-ALC) decision rule:

$$\Pr(correct) = \frac{\exp\left(\frac{a_+}{\tau}\right)}{\exp\left(\frac{a_+}{\tau}\right) + \exp\left(\frac{a_-}{\tau}\right)} \qquad \textit{Eq. S1.1}$$

Where, $a_+$ denotes the activity level of the target, $a_-$ denotes the activity level of the non-target alternative, and $\tau$ is a constant temperature parameter fixed at a value of 2. Unlike the cross-entropy measure, 2-ALC performance allowed us to examine whether the activity of the target output unit was greater than the activity of the concurrently presented non-target (which tended to be activated by its corresponding input).
