## Supplementary material for "Hippocampal and medial prefrontal cortices encode structural task representations following progressive and interleaved training schedules": S2 Text

### Supplementary results for the representational similarity analysis

Here we report significant effects from the representational similarity analyses that did not involve interactions with transitive distance and so are not directly related to our a priori hypotheses. These results therefore reflect changes in representational similarity across all wall textures that only vary by inferential accuracy, transitive slope, or experimental condition. Such effects may suggest that a brain region is involved in transitive inference, yet their precise contributions cannot be inferred from these results alone. This is because there is no unambiguous relationship between uniform changes in pattern similarity across all wall textures, and theoretical accounts of how representational structure may facilitate inference.

#### Within-hierarchy RSA

##### Correlations with inferential accuracy

In both the left entorhinal cortex and left superior MPFC, we detected a main effect of inference accuracy indicating greater pattern similarity between stimulus representations in participants who were more proficient at the inference task, $t(651) = 3.85$, $p < .001$, and $t(650) = 2.95$, $p = .003$ (respectively). These effects were principally driven by accuracy-similarity correlations in the interleaved training condition ($t = 4.51$, & $t = 3.14$) since the effect was non-significant for progressive learners ($t = 1.04$, & $t = 1.34$). Nonetheless, the accuracy by group interaction did not reach our statistical threshold in either ROI; $t(651) = 2.37$, $p = .018$, and $t(650) = 0.89$, $p = .379$, for the left entorhinal cortex and MPFC, respectively. These results suggest that representations in the left entorhinal and medial prefrontal cortices may be involved in successful inference following interleaved training.

##### Correlations with transitive slope

The right inferior MPFC exhibited a main effect of transitive slope such that pattern similarity between wall texture representations was lowest when participants' behavioural performance was most indicative of encoding-based generalisations (i.e., a negative association), $t(654) = 3.29$, $p = .001$. The right superior MPFC also showed a negative correlation between transitive slope and pattern similarity, yet this effect was limited to the interleaved learners, $t = 3.82$. Progressive learners showed the opposing relationship such that behavioural responses strongly indicative of encoding-based generalisations corresponded to higher levels of pattern similarity, $t = 2.48$. This resulted in a training method by transitive slope interaction, $t(652) = 4.50$, $p < .001$. In the left superior MPFC, we detected a significant interaction between training session and transitive slope, $t(650) = 4.38$, $p < .001$. This indicated that pattern similarity was highest in the remote condition when participant's behavioural responses were most consistent with encoding-based generalisations.

##### Categorical effects of condition

Both the left and right superior MPFC showed a significant training method by session interaction, $t(650) = 4.29$, $p < .001$, and $t(652) = 6.34$, $p < .001$ (respectively). These effects indicated that pattern similarity estimates were comparable across all learners in the recent condition yet differed from this level in the remote condition. In the interleaved learners, BOLD responses between remotely trained stimuli were less similar compared with the responses between the recently learnt stimuli. However, the reverse was true for the progressive learners; relative to the recent condition, pattern similarity estimates increased in the remote condition.

#### Across-hierarchy RSA

In a final set of mixed-effects regression models, we examined whether any ROIs exhibited BOLD representations that encoded the structure of the transitive hierarchy in a way that generalised across the transitive chains learnt on each day of training (i.e., across recent and remote conditions). Such representations are predicted by encoding-based models of generalisation, notably, the Tolman Eichenbaum Machine [7]. This analysis involved estimating the similarity between wall-texture representations from different days of training and identifying across-hierarchy distance effects. Similar to above, we tested whether such distance effects were modulated by 3 factors of interest: *1)* training method (interleaved vs progressive), *2)* inference accuracy, and *3)* transitive slope.

None of our regions of interest produced significant across-hierarchy distance effects. However, the left hippocampus, and left and right superior MPFC showed strong correlations between across-hierarchy pattern similarity and our behavioural measures of performance. In the left hippocampus, we observed a significant interaction between training method and transitive slope such that behavioural performance strongly suggestive of encoding-based generalisations was positively associated with increased similarity estimates in the progressive learning condition ($t = 2.63$), but negatively associated with similarity in the interleaved condition ($t = -2.03$); interaction effect: $t(835) = 3.31$, $p < .001$.

Results in the left and right superior MPFC were very similar. Both regions produced a training method by inference accuracy interaction, $t(835) = 4.29$, $p < .001$ and $t(835) = 3.22$, $p = .001$ (respectively). These effects indicated that accuracy scores were positively correlated with representational similarity in the progressive learners, but negatively correlated with similarity scores in the interleaved learners. Both regions also produced a main effect of transitive slope such that participants who produced behavioural responses most indicative of encoding-based generalisations had the highest levels of representational similarity, $t(835) = 3.91$, $p < .001$ and $t(835) = 4.29$, $p < .001$ (respectively).
