## Supplementary material for "Hippocampal and medial prefrontal cortices encode structural task representations following progressive and interleaved training schedules": S3 Text

### Supplementary analyses of the medial prefrontal cortex

To supplement our a priori analyses in the MPFC, we re-ran all univariate and multivariate analyses on MPFC regions of interest derived from a finer parcellation of the cortex – the Schaefer et al 17-network, 400-region parcellation (see Methods). Specifically, we selected MPFC ROIs from this parcellation provided they had at least a 10% volume overlap with our a priori ROIs. This resulted 18 smaller MPFC masks: 4 from the left inferior MPFC, 4 from the left superior MPFC, 6 from the right inferior MPFC, and 4 from the right superior MPFC. We excluded one of these ROIs in the right inferior MPFC as, in several participants, it yielded highly similar BOLD responses with little variability, indicating poor signal levels. This left us with 17 MPFC ROIs and so we only report effects that survive a Bonferroni correction across these regions.

#### Univariate BOLD effects

As in our principal analysis, subregions of the left and right superior MPFC produced BOLD activity associated with behavioural performance. However, these effects were not specific to, or enhanced by, novel inferences as would be expected under retrieval-based models of memory generalisation. Instead, these subregions tended to produce less BOLD deactivations (relative to the implicit baseline) whenever behavioural performance was high, regardless of the trial type (premise vs inferred), training method (interleaved vs progressive), or training session (recent vs remote).

Specifically, a subregion in the right superior MPFC (#369 in the Schaefer atlas) produced a significant 3-way interaction between training session, trial type, and inference accuracy, $t(784) = 4.68$, $p < .001$. This highted that behavioural performance positively correlated with BOLD activity on inference trials in the interleaved learning condition, but higher overall in all other conditions (where performance was close to ceiling). Similarly, although the same 3-way interaction was not significant, $t(786) = 2.32$, $p = .021$, a subregion in the left superior MPFC (#164) revealed an effect of trial type indicating more BOLD deactivation on inference trials (which were associated with lower levels of behavioural performance), $t(786) = 3.96$, $p < .001$.

Unlike in the main text, this supplementary analysis revealed some univariate BOLD effects associated with training session. A subregion in the right inferior MPFC (#314) showed a main effect of session, $t(314) = 3.96$, $p < .001$. This was principally driven by less deactivation on premise trials in the recent condition, although the trial type by session interaction did not meet our significance threshold, $t(786) = 2.18$, $p = .029$. A different subregion of the MPFC (#313) showed a session by transitive slope interaction, $t(786) = 3.14$, $p = .002$. This indicated that transitive slopes were negatively correlated with BOLD activity in the recent condition but positively correlated with BOLD activity in the remote condition, t-values: -2.22 and 2.64, respectively. These effects were not predicated a priori however they suggest that the MPFC may be differentially engaged when behavioural responses depend on information that was learnt at different times.

#### Representational similarity analyses

As in the main text, here we only present multivariate effects involving distance effects that were predicted a priori.

Left superior MPFC

A subregion of the left superior MPFC (#163), showed a significant main effect of distance, $t(652) = 3.59$, $p < .001$, that was superseded by a distance by session interaction, $t(652) = 4.87$, $p < .001$. Similar our main results therefore, this indicated that transitive distance was negatively correlated with representational similarity in the recent, but not the remote, condition. This subregion also produced a distance by transitive slope interaction, $t(652) = 3.66$, $p < .001$, that was superseded by a 3-way interaction between training session distance, and transitive slope, $t(652) = 3.82$, $p < .001$. These latter effects indicated that the distance effects we observed were most pronounced for participants who produced transitive slopes around zero (a behavioural pattern that is less indicative of encoding-based generalisation).

A different subregion in the left superior MPFC (#161), also yielded a significant distance by transitive slope interaction, $t(653) = 4.39$, $p < .001$. In contrast to the above, this effect suggesting that distance effects were strongest for participants who produced larger (more positive) transitive slopes (most typical of encoding-based generalisations).

Right superior MPFC

Consisting with the results of our principal analysis, subregions of the right superior MPFC produced distance effects in the recent condition that were most evident when inference accuracy was relatively low. In one of these subregions (#368), this manifested as a 3-way interaction between distance, session, and inference performance, $t(653) = 3.20$, $p = .001$. Another subregion exhibited the same pattern (#369), yet only a 3-way interaction between distance, inference accuracy, and training session survived our significance threshold, $t(652) = 3.02$, $p = .003$. This indicated the distance effect was most pronounced in interleaved learners who did not achieve high levels of behavioural performance.

Again, replicating an result from the principal analysis, a third subregion of the right superior MPRC (#371) produced a 3-way interaction between distance, session and transitive slope, $t(650) = 3.017$, $p = .001$. This highlighted that the expected distance effects were only expressed in the remote condition when participants' behavioural data was heavily indicative of encoding-based generalisations. Finally, a fourth subregion of the right superior MPFC (#370) produced an interaction between distance, training method, and inference accuracy, $t(651) = 3.15$, $p = .002$. Consistent with our a priori predictions, this indicated that distance effects were strongest in progressive learners who achieved high levels of inference performance.
