## Supplementary material for "Hippocampal and medial prefrontal cortices encode structural task representations following progressive and interleaved training schedules": S1 Table

| | Interleaved training | | | Progressive training | |
|---|---|---|---|---|---|
| | *Recent* | *Remote* | | *Recent* | *Remote* |
| AND model | -.004<br>(.104)<br>[-.191, .219] | .067<br>(.092)<br>[-.120, .239] | | .096<br>(.105)<br>[-.138, .277] | -.230<br>(.074)<br>[-.364, -.070] |
| OR model | .462<br>(.107)<br>[.223, .647] | .545<br>(.127)<br>[.178, .731] | | .543<br>(.074)<br>[.354, .658] | .317<br>(.117)<br>[.043, .514] |
