## Supplementary figures and images for "Hippocampal and medial prefrontal cortices encode structural task representations following progressive and interleaved training schedules"

### S1 Figure

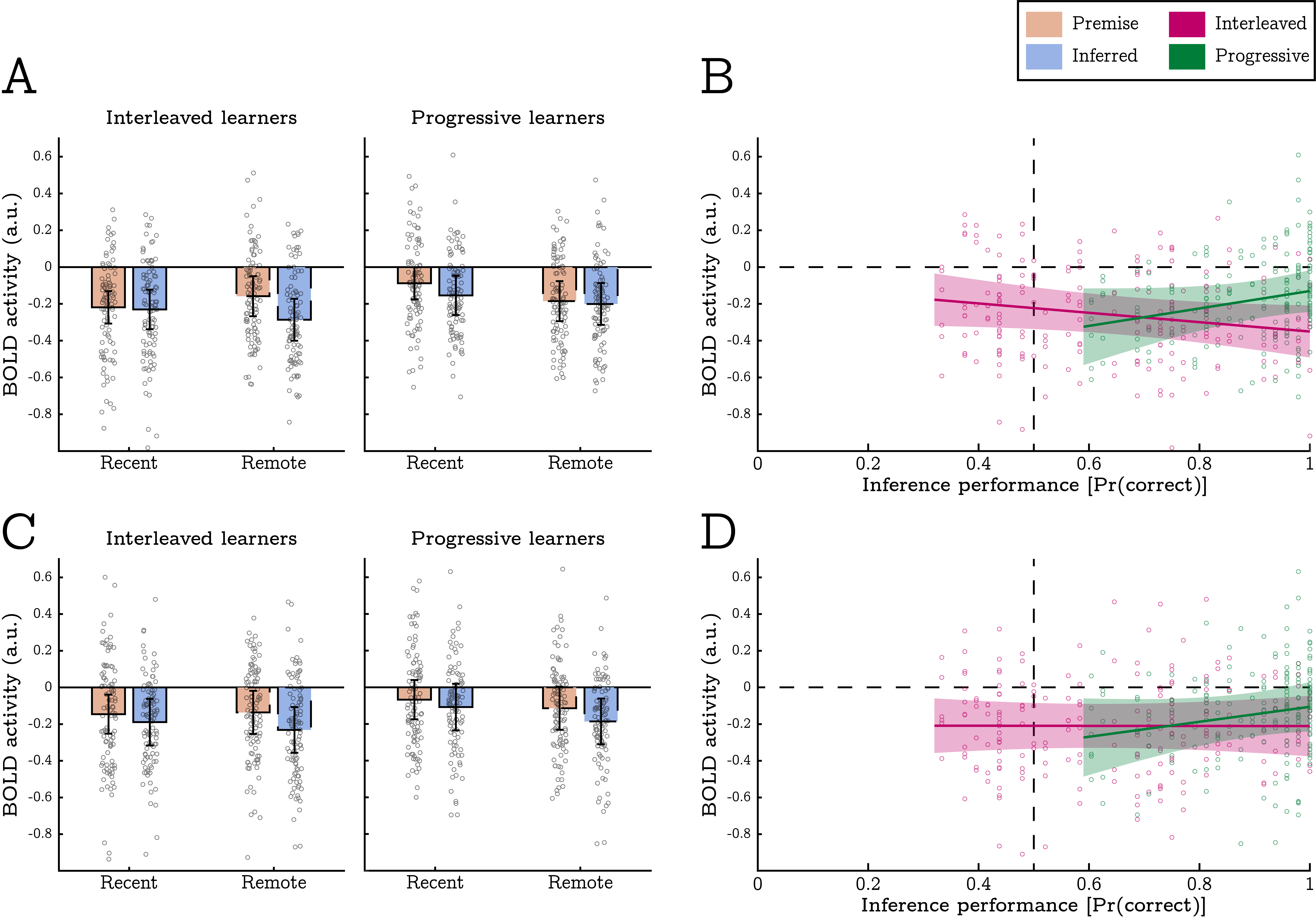

### S2 Figure

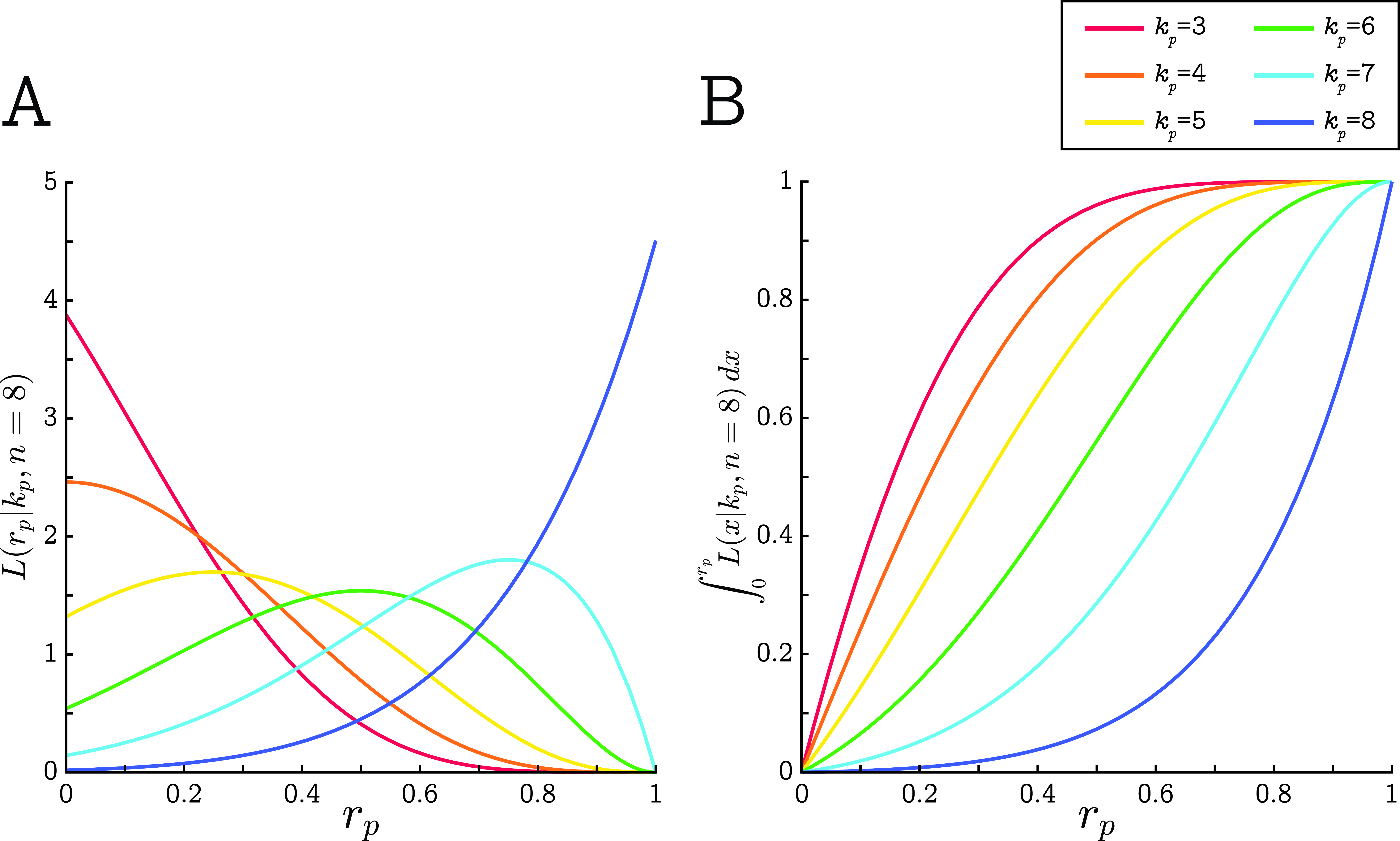

### S3 Figure

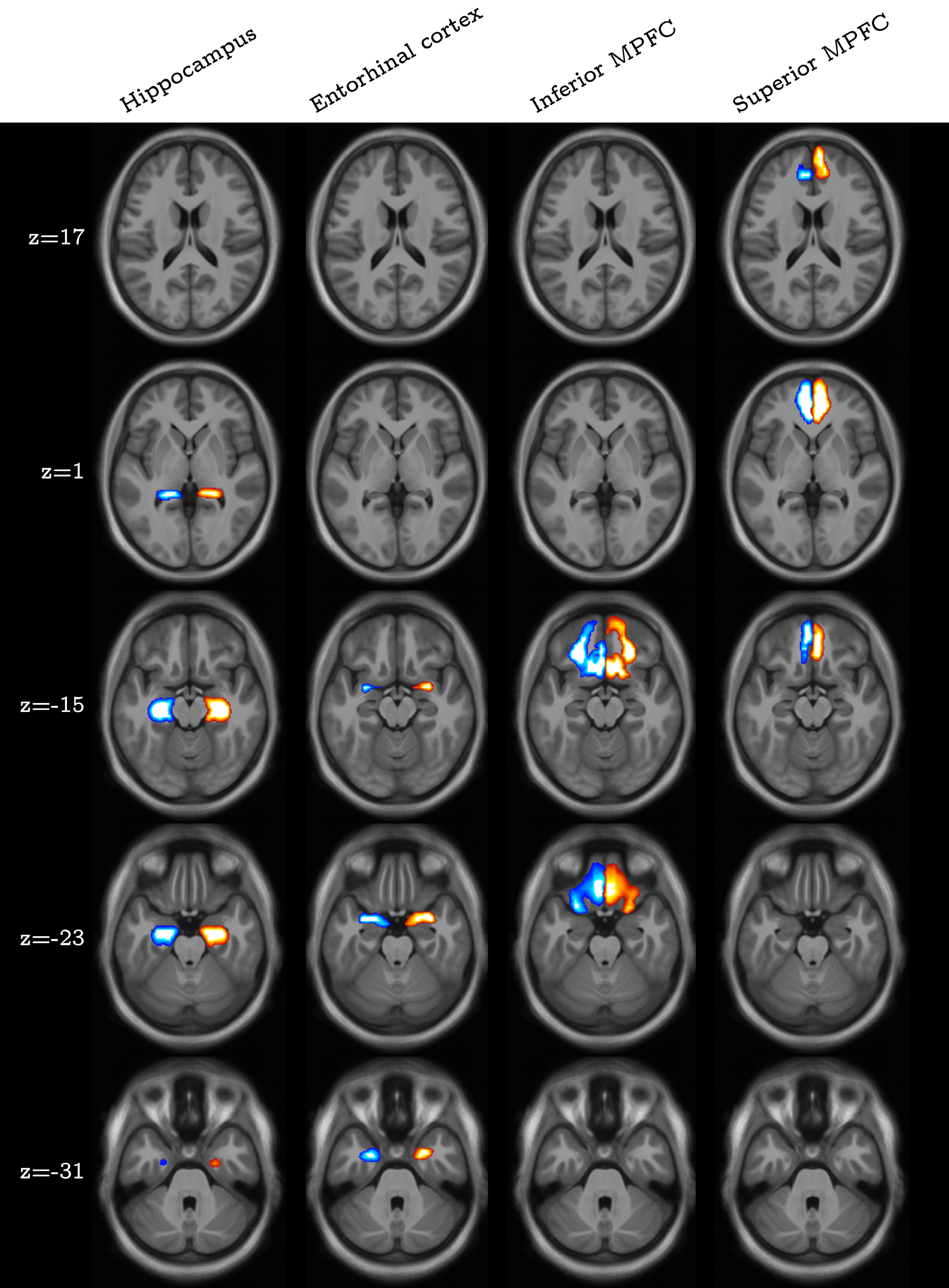
